# The landscape of gene fusions in hepatocellular carcinoma

**DOI:** 10.1101/055376

**Authors:** Chengpei Zhu, Yanling Lv, Liangcai Wu, Jinxia Guan, Xue Bai, Jianzhen Lin, Tingting Liu, Zhang Haohai, Wang Anqiang, Xie Yuan, Wan Xueshuai, Zheng Yongchang, Yang Xiaobo, Miao Ruoyu, C. Robson Simon, Sang Xinting, Chenghai Xue, Haitao Zhao

**Author notes:** Corresponding authors. **Addresses:** Department of Liver Surgery, Peking Union Medical College Hospital, Chinese Academy of Medical Sciences and Peking Union Medical College (CAMS & PUMC), 1 Shuaifuyuan, Wangfujing, Beijing 100730, China.; (X. Sang). My Health Gene Technology Co., Ltd.,Tianjin, China. (C. Xue). Department of Liver Surgery, Peking Union Medical College Hospital, Chinese Academy of Medical Sciences and Peking Union Medical College (CAMS & PUMC), 1 Shuaifuyuan, Wangfujing, Beijing 100730, China.; (H. Zhao). **E-mail addresses:** (X. Sang), (C. Xue), (H. Zhao). These authors contributed equally to this work. These authors share senior co-authorship.

## Abstract

Most hepatocellular carcinoma (HCC) patients are diagnosed at advanced stages and suffer limited treatment options. Challenges in early stage diagnosis may be due to the genetic complexity of HCC. Gene fusion plays a critical function in tumorigenesis and cancer progression in multiple cancers, yet the identities of fusion genes as potential diagnostic markers in HCC have not been investigated.Paired-end RNA sequencing was performed on noncancerous and cancerous lesions in two representative HBV-HCC patients. Potential fusion genes were identified by STAR-Fusion in STAR software and validated by four publicly available RNA-seq datasets. Fourteen pairs of frozen HBV-related HCC samples and adjacent non-tumor liver tissues were examined by RT-PCR analysis for gene fusion expression.We identified 2,354 different gene fusions in the two HBV-HCC patients. Validation analysis against the four RNA-seq datasets revealed only 1.8% (43/2,354) as recurrent fusions that were supported by public datasets. Comparison with four fusion databases demonstrated that three (HLA-DPB2-HLA-DRB1, CDH23-HLA-DPB1, and C15orf57-CBX3) out of 43 recurrent gene fusions were annotated as disease-related fusion events. Nineteen were novel recurrent fusions not previously annotated to diseases, including DCUN1D3-GSG1L and SERPINA5-SERPINA9. RT-PCR and Sanger sequencing of 14 pairs of HBV-related HCC samples confirmed expression of six of the new fusions, including RP11-476K15.1-CTD-2015H3.2.Our study provides new insights into gene fusions in HCC and could contribute to the development of anti-HCC therapy. RP11–476K15.1-CTD–2015H3.2 may serve as a new therapeutic biomarker in HCC.

## Introduction

Hepatocellular carcinoma (HCC) is the third leading cause of cancer-related death worldwide (Zhou et al. 2016). Even though advanced early surveillance technology has improved the life of patients diagnosed at an early stage, most patients are diagnosed with late stage of HCC. Furthermore, HCC patients do not show improved long-term disease-free survival or overall survival after surgical resection and auxiliary medication treatment. (Ye et al. 2003; Kamiyama et al. 2009; Eggert et al. 2013; Llovet et al. 2015). One of the main reasons may lie in the complexity of the genetic background of HCC (Llovet et al. 2015).

Fortunately, recent advances in high throughput sequencing technology have helped provide deeper insights into the genomic and transcriptome landscape of cancer. Using genomic sequencing, researchers have identified large numbers of point mutations, insertions, and deletions as well as chromosome rearrangements in different types of cancers (Gerlinger et al. 2012; Miao et al. 2014; Shibata and Aburatani 2014; Xue et al. 2016). RNA-Seq has become another powerful tool to explore the functional role of these genomic alterations.

Previous studies have demonstrated that fusion genes play an important role in tumorigenesis and cancer progression (Mitelman et al. 2007; Soda et al. 2007) and represent one of the most promising therapeutic targets in human malignancy (Rutkowski et al. 2010; Cortes et al. 2012; Kazandjian et al. 2014; Shaw et al. 2014). The first fusion gene, Philadelphia chromosome, was discovered in 1960 and was approved as the therapeutic biomarker of chronic myeloid leukemia in 2001(Nowell 1960; Topaly et al. 2001; Cohen et al. 2002). According to the FusionCancer and Mitelman Database, nearly 10,000 fusion genes have since been detected (Wang et al. 2015). In addition, several highly recurrent fusion genes in specific tumor types have been well characterized. For example, Soda et al. showed that nearly 6.7% of non-small-cell lung cancer (NSCLC) patients carry the EML4-ALK fusion (Soda et al. 2007). Approximately 55% of prostate cancer show the presence of ERG fusion (Hessels and Schalken 2013). The DNAJB1-PRKACA fusion was found in 100% of fibrolamellar HCC (FL-HCC) (15/15) (Honeyman et al. 2014).

The identity of fusion genes and the recurrent fusion events in HCC have not been comprehensively investigated. A previous study re-analyzed RNA-Seq data of normal liver tissue and HepG2 cells from the National Center for Biotechnology Information Sequence Read Archive database and identified 46 fusion genes (Lin et al. 2014a). Another study only detected five fusion genes from 11 HCC tissues and 11 paired portal vein tumor thrombus tissues (Zhang et al. 2015). Owing to the limited numbers of samples and different analysis strategies, the studies did not identify recurrent fusion genes.

Here we used RNA-seq data of multiple lesions of two HCC patients to explore gene fusions in HCC and validate potential fusions using publicly available RNA-seq datasets of HCC. Our efforts unveiled several novel and recurrent fusions in HCC, suggesting their potential as diagnostic markers or molecular therapeutic targets.

## Results

To gain insight into fusion events in HCC, we performed paired-end RNA sequencing on multiple lesions of two Chinese HCC patients (PI and PII), including adjacent noncancerous liver (PI-N), primary HCC (PI-P), intrahepatic metastases (PI-M), and portal vein tumor thrombus (PI-V) from Patient I, and noncancerous liver (PII-N) and two distant HCCs located in the left (PII-L) and right lobes (PII-R) from Patient II. We obtained ~7,152 million reads (Supplementary Table 1), which is sufficient for transcriptome analysis. We used STAR software(Dobin et al. 2013) to align reads to the hg19 reference genome. An average of approximately 86 million reads per sample was uniquely mapped to the reference genome (Supplementary Table 1). Combing all seven liver samples, we detected about 19,000 annotated genes with a read count more than 10 in at least two samples. We found that 80% (15,302) of expressed genes were protein-coding genes. Interestingly, we also found that 14% (2,651) of expressed genes were long noncoding RNAs (lncRNAs) and 5% were pseudogenes annotated in GENCODE (Supplementary Figure 1A).

To further explore the differences of the transcriptome in the two patients, we merged all samples and found no obvious differences between the transcriptomes. However, in both patients, the HCC tumor samples expressed more genes compared with adjacent noncancerous liver samples, suggesting that HCC tumor samples presented a more complex transcriptome (Supplementary Figure 1B and C). In addition, we found more lncRNAs expressed in the HCC tumor samples in both patients than adjacent noncancerous liver samples, indicating that lncRNAs may play key roles in the tumorigenesis of HCC (Supplementary Figure 1B and C).

## Landscape of fusion events in HCC

Transcriptome analysis has been widely used to identify gene fusions in human cancers (Bao et al. 2014; Stransky et al. 2014; Yoshihara et al. 2015). We used paired-end reads whose two segments mapped to distinct genes to determine gene fusions by STAR using the parameters suggested by Nicolas et al.(Stransky et al. 2014) A total of 2,354 different gene fusions with junction read count more than 3 (Supplementary Figure 2A) was identified in the total combined samples. HCC samples possessed more fusion events and involved fusion genes than adjacent normal tissues (Figure 1A). Furthermore, the number of gene fusions gradually increased across the PI_N, PI_P, PI_V and PI_M samples (Figure 1A), which is consistent with intrahepatic metastasis process. Additionally, PI_M showed the most gene fusion events and most involved fusion genes (Supplementary Figure 2B). The right lesion of patient PII showed 241 fusion events, which is more than 193 of left lesion.

**Figure 1.**
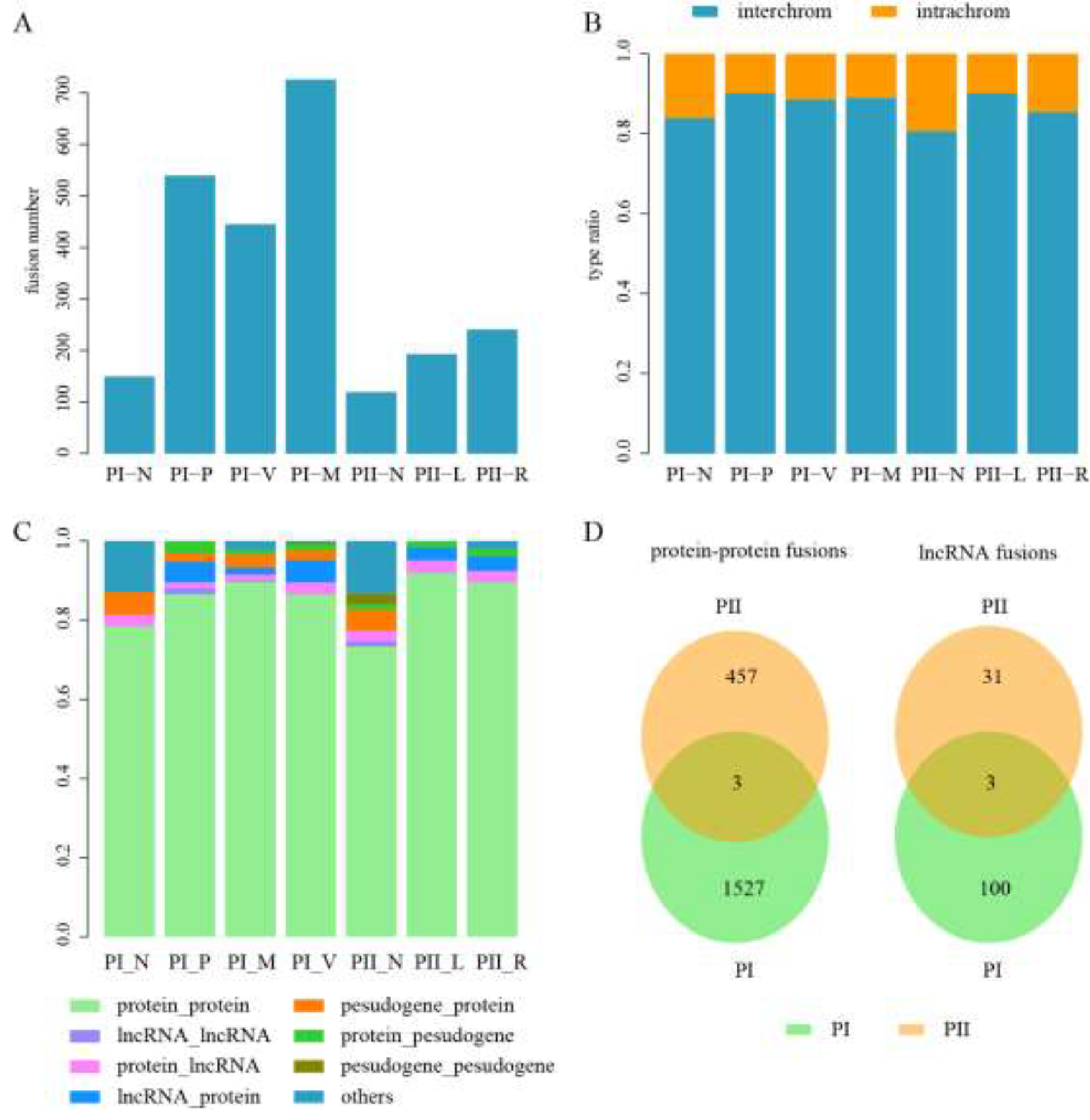
The landscape of fusion events in Hepatocellular carcinoma. (A) The number of fusion events identified in each sample. (B) The ratio of fusion events between different chromosomes and in the same chromosome. (C) The type of two fusion genes involving fusion events. (D). The overlap of two types of fusions between PI and PII, the type of protein-protein fusion represents the fusion between two distinct protein-coding genes and lncRNA fusions represting fusions involved lncRNAs.

By analyzing the genome position of fusion genes, we found that more than 85% fusion events were between two different chromosomes (Figure 1B). We next classified fusion events into eight types according to GENCODE: protein-protein, lncRNA-lncRNA, protein-lncRNA, lncRNA-protein, pseudogene-protein, protein-pseudogene, pseudogene-pseudogene and others. As expected, most fusion events (on average, 85%) were between protein-coding genes in all samples (Figure 1C). The number of fusion types in adjacent normal tissues was less than those in HCC samples, suggesting that more complex fusions were involved in HCC. Moreover, the proportions of distinct fusion types were different across all samples (Figure 1C), and the proportion of protein-ncRNA fusion events in PI was higher than PII. Through comparing the fusion types of protein-protein and lncRNA fusions between PI and PII, we found that the overlap were both small indicating large differences of fusion landscape between two patients.

Interestingly, we found that a few genes can fuse to more than one partner genes (Supplementary figure 3). For example, NDRG1 was a suppressor gene, but it occurred in CCNK––NDRG1 and NDRG1––DUSP3 fusions in PI-V! In summary, there were many fusion events in Hepatocellular carcinoma. Especially there were some fusions involved in ncRNAs (such as lncRNAs and pseudogenes) highlighting the importance of ncRNAs in the development of HCC.

## Recurrent fusion events in HCC

Recurrent fusion events contribute to the development of human cancers and serve as biomarkers or therapy targets of some cancers (Elefanty et al. 1990; Bao et al. 2014; Dhanasekaran et al. 2014). We next examined potential recurrent gene fusions contributing to the pathogenesis of HCC. By combining all gene fusions, we found that majority of gene fusions only occurred in individual samples (Supplementary Figure 4A), and only a few recurrent fusions were present in more than two samples. Of 2,354 fusions, we identified 20 (0.9%) recurrent fusions appearing in more than two samples. As shown in Figure 2A, the recurrent fusions could well classify all samples into two patients. Moreover, except for a few recurrent cases, most fusion genes appeared only once in the seven samples (Figure 2C).

**Figure 2.**
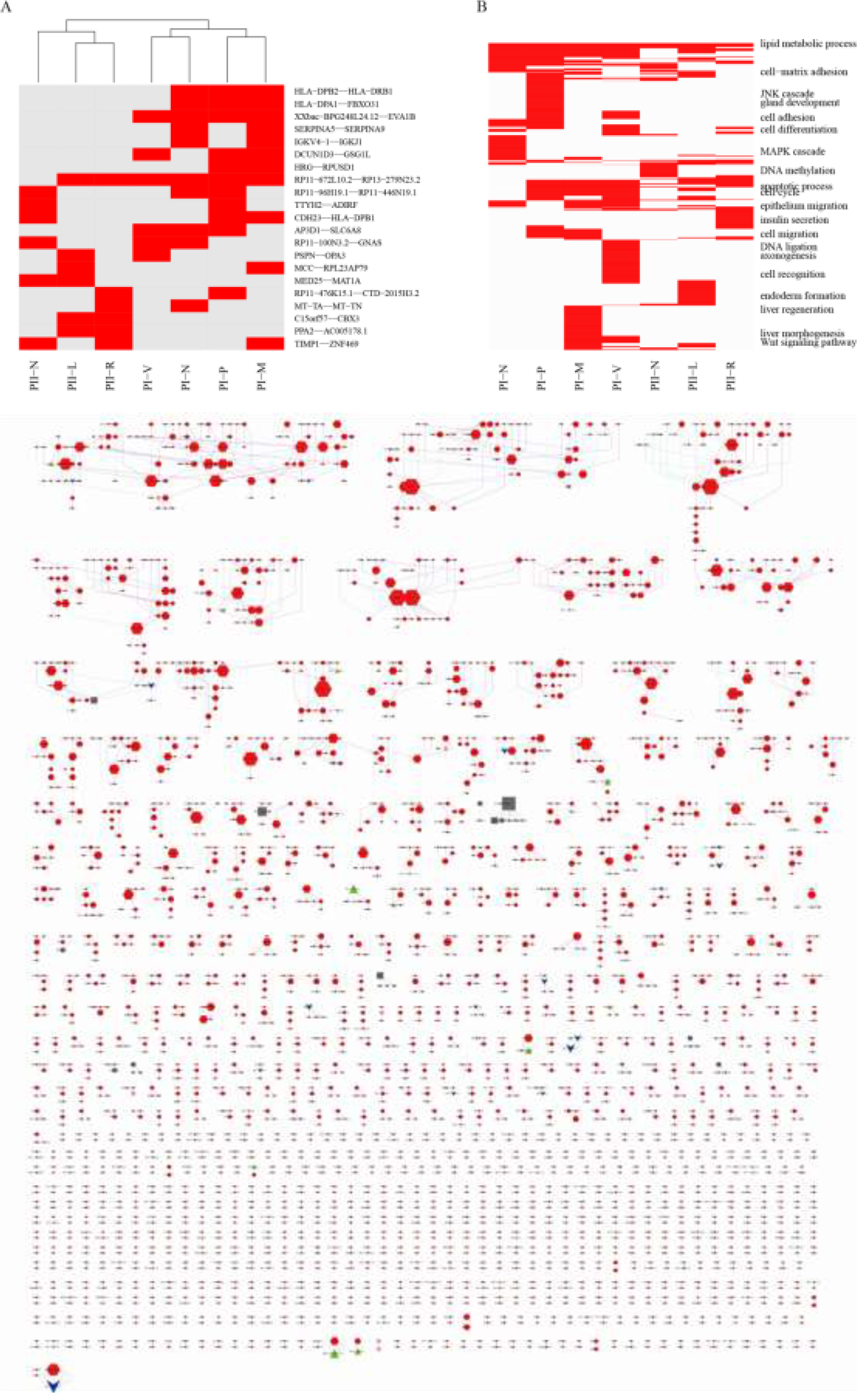
(A) The clusters of fusions appear at least two samples. (B) The enriched functions for all fusion
genes in each sample. (C) Fusion events in 7 samples. 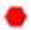 represents protein-coding genes; 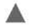 represents pseudogenes; 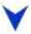 represents lncRNA genes; 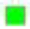 represents other genes;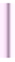 represents fusion event occurred in tumor tissue; 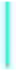 represents fusion event occurred in adjacent non-tumor tissue.

To further explore the functional difference of gene fusions across HCC samples and adjacent normal tissues, we performed functional enrichment analysis using all genes involved in fusion events in each sample. The genes were commonly enriched in several biological processes, such as lipid metabolic process and oxidation-reduction process (Figure 2A and B, Supplementary Figure 4B). HCC samples of PI were enriched in the same biological process, such as cell cycle and apoptotic process. Notably, the majority of enriched functions were only enriched in one sample, suggesting that each sample generated fusion events with specific consequences. For example, fusion genes in PI-P were involved in cancer-related functions and pathways, such as gland development, JNK cascade and p53 signaling pathway. However, fusion genes in PI-M were enriched in migration-related functions, such as cell migration function and focal adhesion pathway. Similarly, the enriched functions of fusion genes in patient PII were also different. Fusion genes in PI-R were involved in Notch signaling pathway regulation (Figure 2B and Supplementary Figure 4B).

Owing to our small sample size, our initial findings may exclude some important recurrent fusions in HCC. Therefore, we collected public available RNA-seq data of HCC to identify reliable recurrent fusions. We obtained four RNA-seq datasets (accession number: GSE65485 (Dong et al. 2015), GSE55758 (Gao et al. 2015), GSE33294 (Chan et al. 2014) and SRP007560 (Lin et al. 2014b); Supplementary Table 2) involved 79 public HCC samples using keywords of “hepatocellular carcinoma”, “HCC”, “liver”, “RNA-seq”, “next generation sequencing”, and “high through output” in GEO database and NCBI SRA database.

To obtain recurrent fusions, we selected 2,354 fusion events with at least three junction reads. In the public datasets, we obtained 24,960 fusion events in total, including 23,795 fusions in GSE65485, 1,132 fusions in GSE55758, 40 fusions in GSE33294, and 92 fusions in SRP007560. We obtained 43 gene fusions that occurred in at least two samples in our HCC samples, or presented in one sample in our data and also occurred in at least one sample in the 79 public samples. The remaining 2,311 gene fusions occurred just once in our seven samples, with no events detected in the public data (Table 1). We speculated whether recurrent gene fusions significantly presented in public samples. We thus grouped the 2,354 fusions into two classes: 20 recurrent fusions present in at least two samples in the seven HCC samples and 2,334 fusions that occurred in one out of seven samples. We found that 15 recurrent fusions presented in at least one of the 79 public samples and five recurrent fusions only were detected in at least two samples of the seven HCC samples. Among the 2,334 fusions, 23 fusions were detected in at least one of 79 public samples. We found that recurrent fusions were significantly supported by public datasets (Fisher’s exact test, p-value < 2.2e–16, Table 1), suggesting that recurrent fusions were likely functional.

**Table 1.**
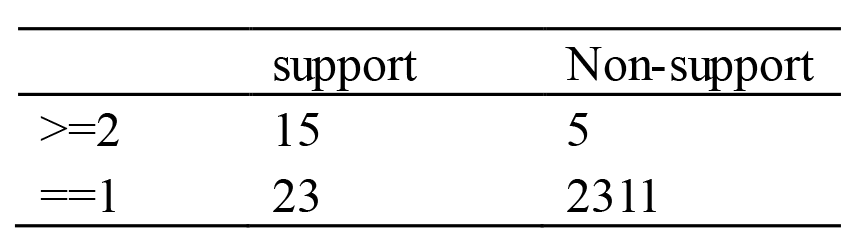
The number of fusions with once fusions and recurrent fusions supported by public HCC samples.

|  | support | Non-support |
| --- | --- | --- |
| $\geq 2$ | 15 | 5 |
| $= 1$ | 23 | 2311 |

## Identification of candidate fusion events associated with HCC

We used four known fusion databases including ChimerDB, FusionCancer, Mitelman and MCL to examine if the identified fusion events were associated with human diseases. In addition, we gathered five sets of tumor-associated fusion events derived from recently published literatures(1Stransky et al. 2014),(Bao et al. 2014),(Dhanasekaran et al. 2014),(Sia et al. 2015),(Torres-Garcia et al. 2014). We examined the distribution of fusion events from the seven HCC samples in the fusion databases and recently published literature. Three (RNF213-SLC26A11, PSPC1-MPHOSPH8 and E2F4-RPL14) of 2,311 fusion events that occurred in only one sample (without any published HCC samples support) were annotated in known fusion databases and recently published literatures. The PSPC1-MPHOSPH8 fusion in PI-M was annotated as t (13; 13) (q12; q12) in the Mitelman database. The E2F4-RPL14 fusion in PI-V was related to Burkitt lymphoma in the FusionCancer database. The RNF213-SLC26A11 fusion event in PII_R was a glioblastomas-related fusion in the study of Zhao et al. and also annotated to chronic myelogenous leukemia in FusionCancer. This indicated that some of the once occurring fusion events may be functional and related to the pathogenesis of HCC.

Three (HLA-DPB2-HLA-DRB1, CDH23-HLA-DPB1, and C15orf57-CBX3) out of 43 recurrent gene fusions were annotated as disease-related fusion events. The fusions HLA-DPB2-HLA-DRB1 and CDH23-HLA-DPB1 were both annotated as a lung cancer fusion in the FusionCancer database. Additionally, 21 out of the remaining 40 recurrent fusion events contained at least one fusion gene that was annotated to participate in at least one disease in the known fusion databases and published literatures. For example, IGLV4-69-IGLJ3 fusion occurred in PI-M, and Daniela et al. demonstrated that LOC9610-IGLJ3 fusion was associated with intrahepatic cholangiocarcinoma. Therefore, these recurrent gene fusions were most likely involved in the tumorigenesis of HCC.

To identify candidate fusion events associated with tumorigenesis of HCC, we integrated known fusion annotations and public available HCC samples. The candidate HCC-associated potential gene fusions should not only be annotated in known fusion databases and recently published literatures, but also occur in as many samples as possible. We calculated the frequency of each recurrent fusion and obtained the ratio of the disease samples compared with the normal liver samples. The fusions showing larger ratio than that of known cancer-related fusions were considered as candidate fusion events associated with tumorigenesis of HCC. The ratios of three known fusions HLA-DPB2-HLA-DRB1, CDH23-HLA-DPB1 and C15orf57-CBX3 were 2, 2.6 and 5, respectively. We selected 17 fusions with ratios more than 2, and 9 fusions that occurred in only cancer samples as candidate fusion events. For example, the fusion C15orf57-CBX3 occurred in PII-L and PII-R. C15orf57-CBX3 was also present in 22.7% (18) public HCC samples and four normal liver samples (Supplementary Table 3). The CBX3 gene encodes a DNA-binding protein that is a component of heterochromatin. CBX3 protein is also recruited to sites of ultraviolet-induced DNA damage and double-strand breaks. C15orf57 is an uncharacterized gene. Furthermore, we found that the breakpoint (hg19 genome location, chr15:40854971; corresponding to GRCh38/hg38 location, chr15:40562772) in the second exon of C15orff57 was located in the CCDC32 conserved domain (pfam14989), suggesting that the domain may be truncated and the protein structure was affected by the fusion. The breakpoint in the first exon of CBX3 (chr7:26241389) is located upstream of the CHROMO and ChSh conserved domains without destroying the domain (Figure 3A and Supplementary Figure 5A). Therefore, the C15orf57 altered domain may join with the complete CBX3 protein to generate a chimeric protein. Moreover, the C15orf57-CBX3 fusion was reported in glioblastoma by Zhao et al. and is associated with cervical cancer, melanoma and Burkitt lymphoma in the FusionCancer database. Thus, the C15orff57-CBX3 fusion may be involved in the development of HCC.

**Figure 3.**
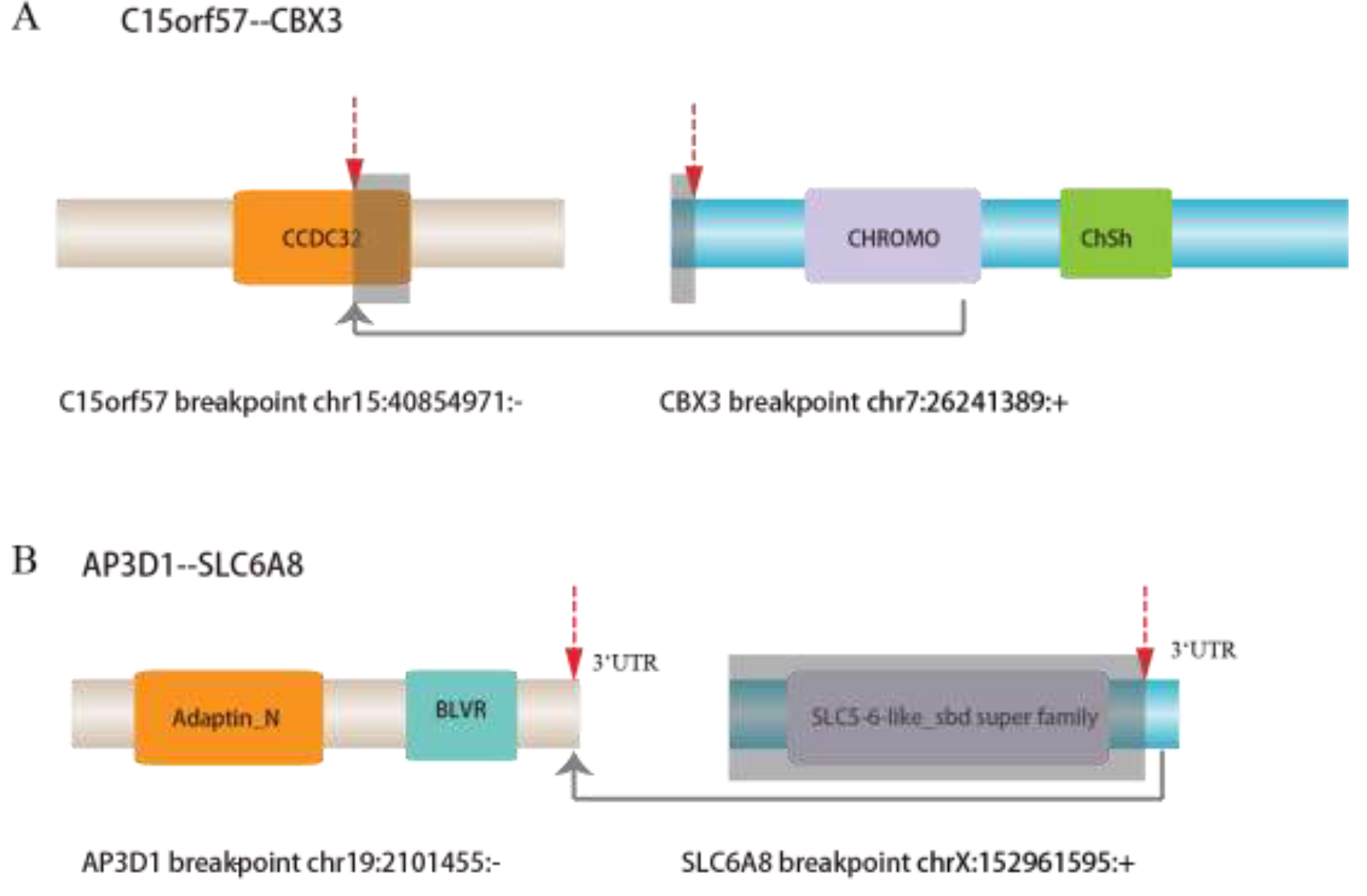
The fusion events involved in known disease related fusion genes. (A) The breakpoint of known disease fusion C15orf57––CBX3. (B) The breakpoint of AP3D1––SLC6A8.

Another fusion, AP3D1-SLC6A8, which occurred in PI-N, PI-P and PI-V with abundant of junction reads, was also present in nine public HCC samples without any supporting normal public liver sample. AP3D1-SLC6A8 showed the highest ratio of disease sample (11 HCC samples vs. 1 adjacent noncancerous liver PI-N, Supplementary Table 4). The AP3D1 gene encodes a protein subunit of the AP3 adaptor-like complex, which is associated with the golgi region, as well as more peripheral structures, suggesting an important role in the translation process(Petrenko et al. 2006). The SLC6A8 gene encodes a plasma membrane protein that transports creatine into and out of cells and is also involved in the ion transport process. Furthermore, we found that the breakpoint (chr19:2101455) in the 3′ UTR of AP3D1 and does not affect protein domains. The breakpoint (chrX: 152961595) of SLC6A8 was also located in the 3′ UTR (Figure 3B and Supplementary Figure 5B). Therefore, the AP3D1-SLC6A8 fusion retains full-length AP3D1 and part of 3′ UTR region of SLC6A8. The fused 3′ UTR of SLC6A8 may influence the regulation of AP3D1.

Moreover, AP3D1 was reported to be fused with the NSUN2 gene in HCC in the FusionCancer database. In addition, AP3D1 was reported to be involved in fusions in cervical cancer, lung cancer and colon cancer in the FusionCancer database, indicating that AP3D1 may be widely involved in fusions in human cancer. SLC6A8 was annotated to participate in melanoma (SLC6A10P-SLC6A8) and glioblastomas (SLC6A8-GABRA3) in FusionCancer. Thus, the AP3D1-SLC6A8 fusion may be associated with tumorigenesis of HCC and should be further examined.

## Novel fusion events associated with HCC

In addition to recurrent gene fusions involving known fusion genes or fusion events, we found 19 novel and recurrent fusions that have not been previously annotated to diseases (Supplementary Table 7). For instance, the recurrent fusion DCUN1D3-GSG1L occurred in PI-P, PI-V and PI-M without any public sample support. Overexpressing DCUN1D3 gene may promote mesenchymal to epithelial-like changes and inhibit colony formation in soft agar(Huang et al. 2014). GSG1L is a component of the inner core of the AMPAR complex, which modifies AMPA receptor gating. Furthermore, both fusion genes harbored the exact same breakpoint in three HCC samples (Figure 4A and Supplementary Table 5). The breakpoint (chr16:20871370) in the third exon of DCUN1D3 is located in the DUF298-conserved domain that binds to cullins and Rbx-1, components of an E3 ubiquitin ligase complex for neddylation. The protein structure is affected by the fusion. The breakpoint in the sixth exon of GSG1L (chr16:27802788) is located downstream of the protein’s conserved domain without affecting any doma ins. Thus, the fusion DCUN1D3-GSG1L may be involved the development of HCC and should be examined.

**Figure 4.**
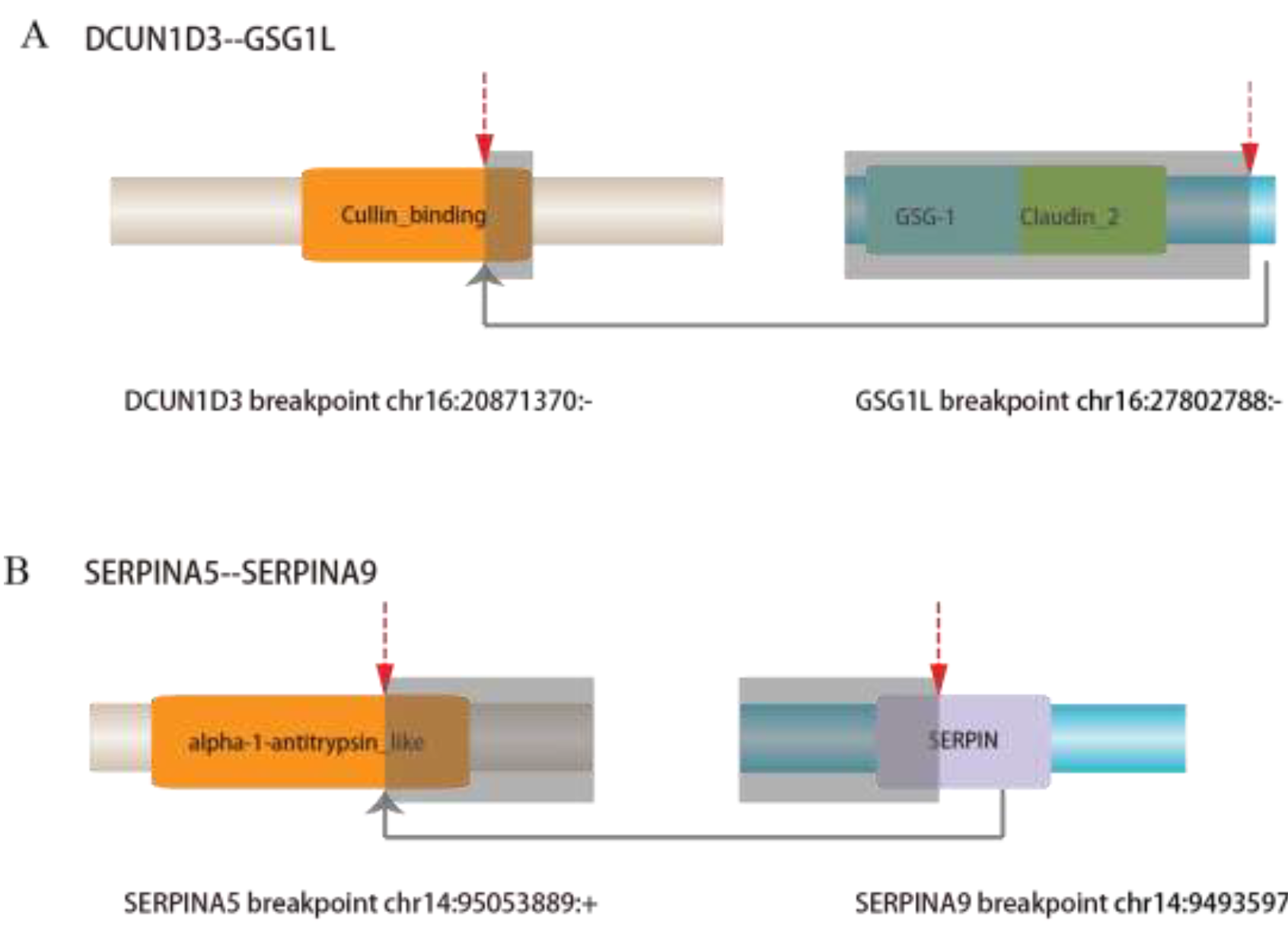
The breakpoint of novel fusion events associated with Hepatocellular carcinoma.

Another novel recurrent fusion SERPINA5-SERPINA9 occurred in PI-N and PI-M, as well as nine public HCC samples (Figure 4B and Supplementary Table 6). SERPINA5-SERPINA9 showed the second highest ratio of disease (10 HCC samples vs. 1 adjacent noncancerous liver PI-N). SERPINA5 is a serpin peptidase inhibitor (Jing et al. 2014) with serine-type endopeptidase inhibitor activity. The breakpoint (chr14:95053889) in SERPINA5 and the breakpoint (chr14:94935978) in SERPINA9 were both located in conserved domains of each protein, suggesting that the domains are truncated in the fusion protein. The fusion can generate a chimeric protein that may be involved in the tumorigenesis of HCC and should be further validated.

## Validating candidate recurrent fusion genes in clinical patients with HCC

We identified 43 recurrent fusion genes for further experimental validation with RT-PCR and Sanger sequencing. And we successfully obtained primer sequences of 26 fusion genes (Supplementary Table 7). The primer sequences are listed in Supplementary Table 8. We selected 11 pairs of HBV-related HCC samples combined with their corresponding adjacent non-tumor liver tissues and 3 multiple lesion patient samples (Patient 2, Patient A and Patient B). Samples collected from Patient 2 included noncancerous liver, and two distant HCCs located in the left and right lobes; in Patient A, noncancerous liver, tumor lobe and portal vein tumor thrombus; and from patient B, noncancerous liver and two distant HCCs. Six fusion genes were validated to exist in clinical samples (Table 2, Figure 5, and Supplementary Figures 6–9).The detailed sequences of these fusion genes are listed in Supplementary Data A. Five of the six validated fusion genes (except for IGLV4-69-IGLJ3) were detected in many clinical samples both in the tumor samples and adjacent noncancerous samples.

**Figure 5.**
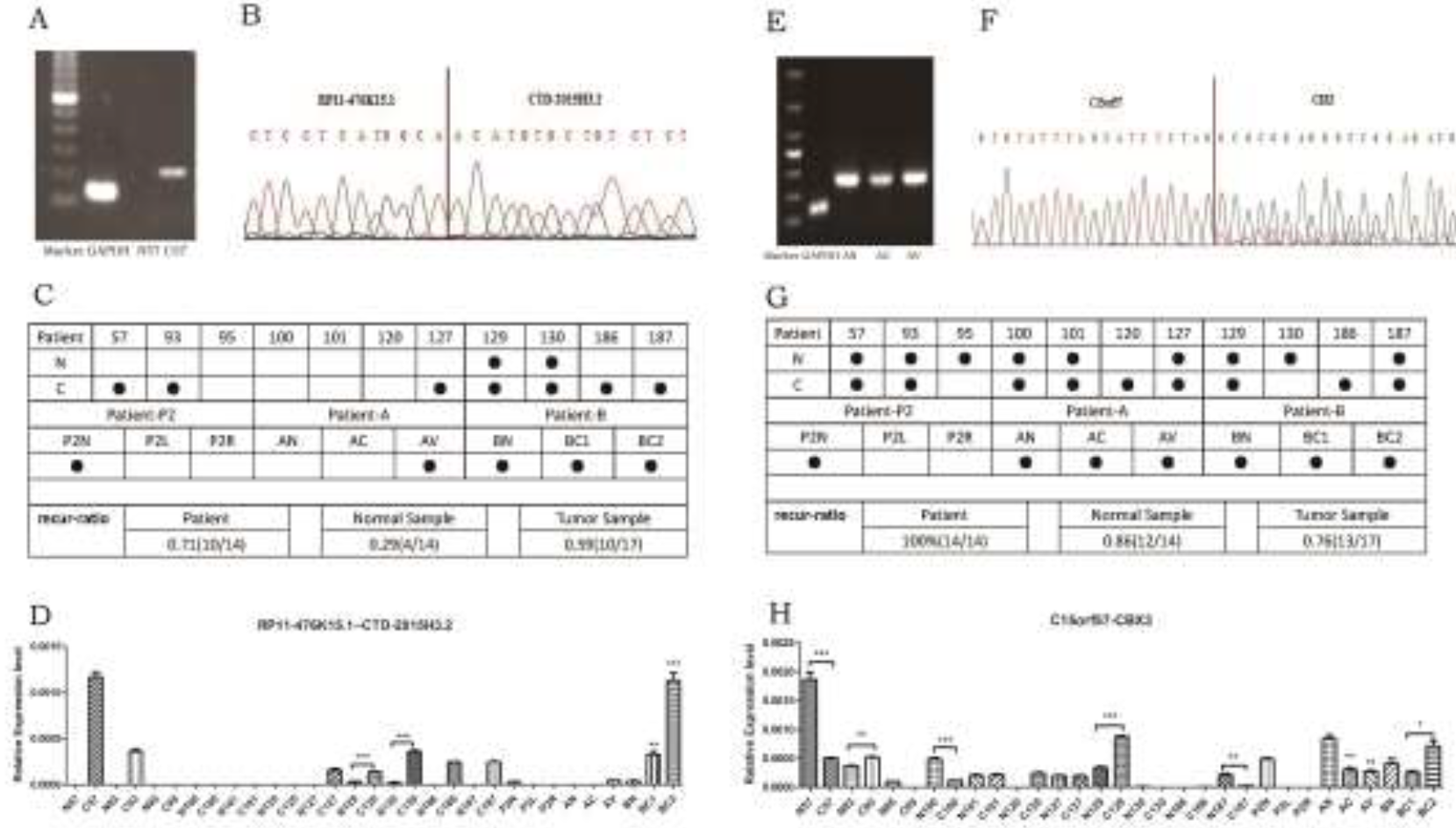
Details of two fusion genes after experimental validation of the fusion transcripts by RT-PCR, Sanger sequencing and Quantitative Real-time PCR. (A-B) The electrophoretic result and sequencing data for RT-PCR product with fusion gene RP11-476K15.1––CTD-2015H3.2. (C) Verified samples for the existence fusion RP11-476K15.1––CTD-2015H3.2. • represents the verified sample. Recur-ratio shows the ratio of the verified sample comparing with the total number in patient, normal (noncancerous) sample and tumor sample, respectively. (D) The relative expression level of fusion gene RP11-476K15.1––CTD-2015H3.2 in HCC samples. (E-H) Demonstrate the result of the fusion gene C15orf57––CBX3, possess similar annotation information with fusion gene RP11-476K15.1––CTD-2015H3.2.

**Table 2.** Details of candidate recurrent fusion genes after experimental validation of the fusion transcripts by RT-PCR and Sanger sequencing in HCC samples.

| fusion_name | chromosome |  | Supported normal samples | Supported tumor samples |
| --- | --- | --- | --- | --- |
| C15orf57--CB<br>X3 | Chr15-chr7 | inter- | 12<br>N57/N93/N95/N100/N101/N<br>127/N129/N130/N187/<br>P2N/AN/BN | 13<br>C57/C93/C100/C101/C120/C127/C<br>129/C186/C187/AC/AV/BC1/BC2 |
| IGLV1-51--IG<br>LL5 | Chr22-chr22 | intra- | 10<br>N93/N95/N101/N120/N129/<br>N130/N187/ P2N/AN/BN | 13<br>C57/C93/C127/C120/C129/C130/C<br>186/C187/ P2L/P2R/AV/BC1/BC2 |
| IGLV4-69--IG<br>LJ3 | Chr22-chr22 | intra- | 3<br>N95/ N100/ P2N | 2<br>AC/AV |
| RP11-100N3.2<br>--GNAS | Chr11-chr20 | inter- | 9<br>N57/N93/N95/N100/N101/N<br>127/N129/N186/ P2N | 8<br>C93/C95/C127/C129/C130/C187/<br>P2R/BC1 |
| RP11-476K15.<br>1--CTD-2015H<br>3.2 | Chr18-chr18 | intra- | 4<br>N129/N130/ P2N/BN | 10<br>C57/C93/C127/C129/C130/C186/C<br>187/ AV/BC1/BC2 |
| XXbac-BPG24<br>8L24.12--EVA<br>1B | Chr6-c<br>hr1 | inter- | 12<br>N57/N93/N95/N100/N101/N<br>120/N127/N129/N130/N186/<br>P2N/AN | 16<br>C57/C93/C95/C101/C120/C127/C1<br>29/C130/C186/C187/P2L/P2R/AC/<br>AV/BC1/BC2 |

We analyzed the relative gene expression in tumor samples and adjacent noncancerous samples using quantitative real-time polymerase chain reaction (qRT-PCR) (Figure 5D, 5H and Supplementary Figure 6–9). Among the 26 candidate recurrent fusion genes, six fusions were confirmed by RT-PCR and Sanger sequencing (Figure 5A, B, E, F and Supplementary Figure 6–9). Though some fusions were detected more frequently in tumor samples compared with adjacent noncancerous samples, many fusions were frequently detected both in clinical tumor and adjacent noncancerous samples. For instance, the newly identified fusion RP11-476K15.1-CTD-2015H3.2 was detected in our HCC samples PI_P and PII_R. For the validated fusions that occurred in tumor and benign samples showed relatively higher expression in tumor samples compared with noncancerous samples (Figure 5A–C). RP11-476K15.1-CTD-2015H3.2 was identified in 71% (10/14) of patients, 29% (4/14) of noncancerous samples, and 59% (10/17) of tumor samples (Figure 5C). These findings suggest that RP11-476K15.1-CTD-2015H3.2 is a novel HCC-related fusion gene that may be a new therapeutic biomarker or therapy target. Another fusion C15orf57-CBX3 was detected and showed a considerable expression level in tumor samples and noncancerous samples, similar to other 4 fusions (Figure 5G, H and Supplementary Figure 6–9). C15orf57-CBX3 was identified in 100% (14/14) of patients, 86% (12/14) of noncancerous samples, and 76% (13/17) of tumor samples (Figure 5G). We suspect that these clinical patients have had a history of hepatitis B virus infection for several years, and have taken place in liver cirrhosis, which is not fully normal liver tissue. In the process of liver cirrhosis, the genome of liver tissue changes dramatically, resulting in fusion events.

## Kinase gene fusion in HCC

Kinase gene fusion plays an important role in tumorigenesis and targeted therapy. However, few recurrent kinase fusion events have been reported. To examine kinase gene fusion in HCC, we selected fusion events with at least one participating kinase gene. Interestingly, unlike the previous study of Stransky et al., we found many kinase genes in fusion events in both HCC and adjacent non-tumor tissues (Figure 6). For example, MAP3K11 fusion was detected in 23.5% (4/17) of adjacent non-tumor tissues. Both BRD4 and NRBP2 were detected in 25.8% (16/62) of HCC tissues. MET fusion was only detected in 4.8% (3/62) and 5.9% (1/17) of HCC tissues and adjacent non-tumor tissues, respectively.

**Figure 6.**
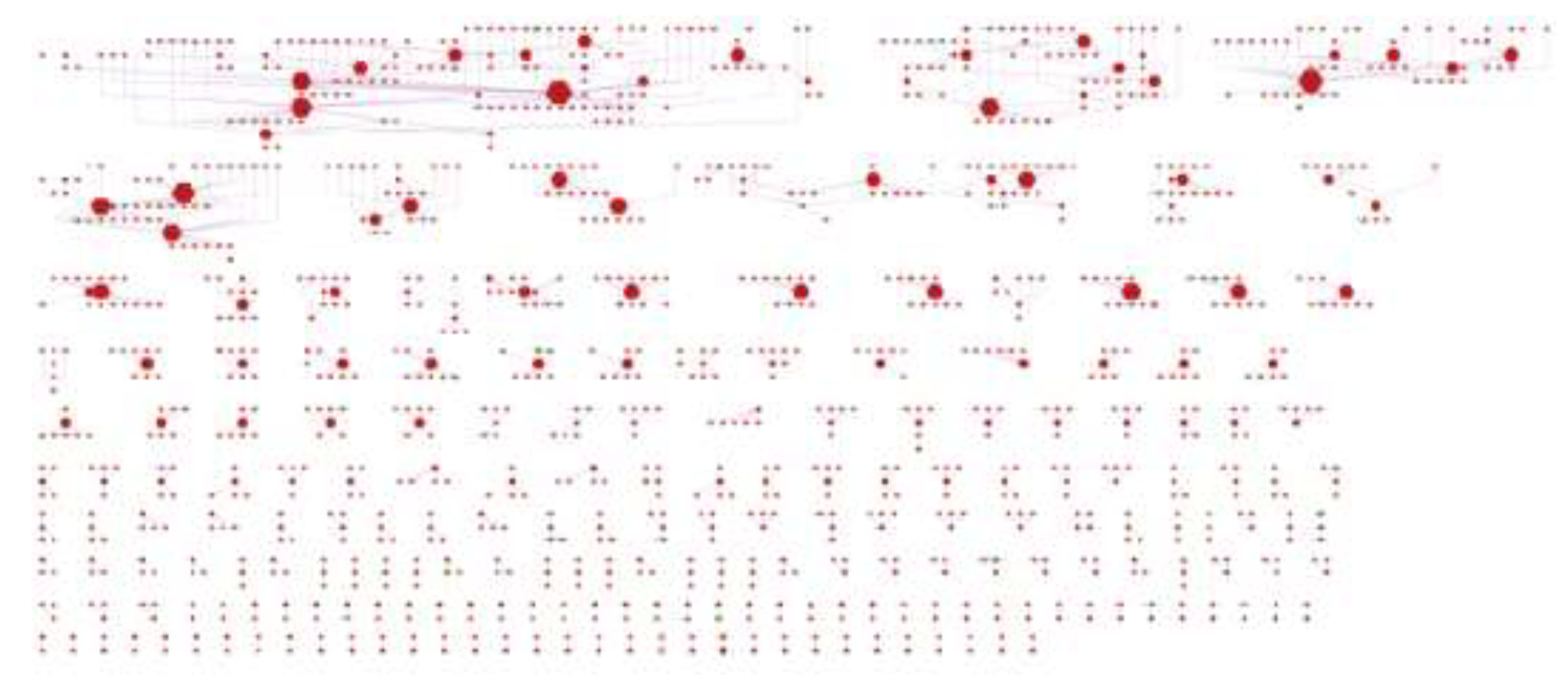
Kinase gene fusions in HCC. 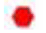 represents kinase gene; 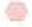 represents ordinary protein-coding gene; 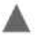 represents pseudogenes; 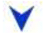 represents IncRNA genes; 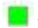 represents other genes; 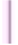 represents fusion event occurred in tumor tissue; 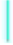 represents fusion event occurred in adjacent non-tumor tissue.

## Discussion

Here, we characterized the landscape of fusions in HCC and identified 43 fusion events and confirmed 26 by RT-PCR and Sanger sequencing. Detection and characterization of fusion genes has been critical in understanding tumorigenesis, anticancer drug screening, and clinical application (Hessels and Schalken 2013; Stransky et al. 2014; Mertens et al. 2015; Yoshihara et al. 2015). However, few fusion events are demonstrated recurrent events. Fortunately, with the development of high throughput sequencing technology as well as bioinformatics algorithms, huge amount of fusion events has been detected(Kim and Salzberg 2011; Li et al. 2013).We detected a total of 2,354 candidate fusion events and only 1.8% (43/2354) of these events were recurrent. Similarly, Yoshihara et al. (Yoshihara et al. 2015) and Stransky et al. (Stransky et al. 2014) reported large numbers of kinase gene fusions in 13 and 20 types of cancer, respectively. However, only 7.4% and 12% were recurrent events.

Several previous studies have focused on fusion events involving protein kinase genes (Stransky et al. 2014; Yoshihara et al. 2015) because of their critical functions in cellular processes and potential targets for anti-cancer drugs. In our analysis, we did not detect protein kinase genes involved in fusion events. This difference may result from the cut-off value in the detecting process. However, some of the identified fusion proteins in our study may have important functionality in HCC. For example, C15orf57-CBX3, a recurrent fusion in 22.7% public HCC samples and 4 normal liver samples, was also a fusion protein in glioblastomas (Bao et al. 2014) and is also annotated to associate with cervical cancer, melanoma and Burkitt lymphoma in the FusionCancer database (Wang et al. 2015). Other fusion proteins, such as AP3D1-SLC6A8, DCUN1D3-GSG1L, and SERPINA5-SERPINA9, may also play an important role in HCC. Further studies should focus on the function of these recurrent fusion proteins.

Noncoding genes also play an important role in human disorders as well as cancer (Guarnerio et al. 2016; Xu et al. 2016). However, to our best knowledge, up until now, few studies have focus on noncoding gene fusion events. Lau et al. reported that HBx-LINE fusion, which functions as an lncRNA, affects β–catenin transitivity and is involved in liver cancer development and progression (Lau et al. 2014). Dong et al. found that HCC patients carrying the HBV-MLL4 fusion have a distinct gene expression profile (Dong et al. 2015). In our analysis, we identified many fusion events involving noncoding genes. Notably, a higher percentage of noncoding gene fusion events were detected in the advanced HCC patient. This supports the idea that noncoding gene fusions play key roles in the progression of cancer.

We also detected many recurrent fusion events in adjacent normal tissue. For example, 27.9% (12/43) of fusion events were detected in only adjacent normal tissues and 30.2% (13/43) recurrent fusion events were observed in both HCC and adjacent normal tissues. Some were highly detected in adjacent normal tissue compared with the paired HCC tissue. Similarly, the TEL-AML1 fusion gene was reported to occur 100 times more frequently in normal individuals than leukemia patients and contributes to initiation of childhood ALL(Mori et al. 2002; Zelent et al. 2004). Thus, fusion events in adjacent normal tissue may serve as biomarkers of hepatitis disease progression into HCC and should be pursued in future experimental study. Successful applied drugs target on BCR-ABL1 fusion in hematological malignancy ALK fusion in NSCLC dramatically ignite the enthusiasm of deep exploration the landscape of gene fusions(Mertens et al. 2015). Moreover, several drugs, such as imatinib and crizotinib, have been approved for targeting gene fusions in human malignant diseases. These success stories support the notion that fusion events represent promising anticancer targets. Our present result provides insight into the landscape of gene fusions in HCC and might pave the way for anti-HCC therapy.

## Materials and Methods

### Patients and clinical samples

Two representative HBV-HCC patients who had tumor resection were selected for next generation sequencing studies. Each patient underwent the same pathologic evaluation on all tumors. Samples were collected from adjacent noncancerous liver (PI-N), primary HCC (PI-P), intrahepatic metastases (PI-M), and portal vein tumor thrombus (PI-V) from Patient I; and noncancerous liver (PII-N) and two distant HCCs located in the left (PII-L) and right lobes (PII-R) from Patient II.

The cDNA libraries were constructed using previous studies (Miao et al. 2014) and sequenced using Illumina HiSeq ^™^ 2000. Finally, 90*2 bp paired-end reads were obtained for each sample. The raw RNA-seq data were deposited at the European Genome-phenome Archive with accession number EGAS00001000372.

### Gene annotations

The reference genome sequence was downloaded from the UCSC database (version hg19, http://genome.ucsc.edu/). The annotation of genes was downloaded from the GENCODE database (http://www.gencodegenes.org/). To identify fusions, we select the v19 version, which includes a comprehensive list of protein-coding genes and lncRNAs. A total of 57,820 genes were obtained, including 20,345 protein-coding genes, 20,345 pseudogenes and 13,870 lncRNAs.

### Analysis of transcriptome data

For raw RNA-seq data, we first assessed the quality of sequencing reads by Fastqc software and then discarded low quality reads with a quality score less than 20 using the trimmomatic tool. Next, for each sample, we aligned the cleaning read to the hg19 reference genome using STAR v2.4.1, which is an ultrafast RNA-seq aligner. For each read, no more than two mismatches were allowed in the alignment process. Reads that were mapped to distinct genes in the reference genome were output into the chimeric-reads file and used for detection of fusion genes.

HT-seq was used for calculated read count for each gene annotated in the EMSEMBL database.

### Identification of fusion genes

We used STAR-Fusion integrated in STAR software to identify potential fusion genes. Reads deposited in the chimeric-reads file indicated putative fusions. STAR-Fusion used reads that aligned to distinct genes to detect candidate fusion genes. To reduce the number of false positive fusion genes, the length of one read aligned to distinct genes was not less than 15bp. Putative fusions between homologous genes were also discarded. Furthermore, we remained fusions with at least three junction reads, which provides direct evidence about a fusion. We also removed the fusions between mitochondria and autosome.

### Functional enrichment analysis

The significance of enrichment analysis of differentially expressed protein-coding genes (DEGs) with GO terms and pathways was determined using a hypergeometric test with FDR correction (FDR ≤ 0.05). In addition, DEGs overlapped at least two genes with the enriched GO terms and pathways.

### Validating experiments for candidate recurrent fusion genes

#### Patients and clinical samples

Eleven pairs of frozen HBV-related HCC samples combined with their corresponding adjacent non-tumor liver tissues and three multiple lesions of patient samples were obtained from HBV-HCC patients who underwent hepatectomy at Peking Union Medical College Hospital (PUMCH). All patients had pathologically confirmed HCC and did not receive any anti-cancer treatment prior to surgery. Fresh tissue samples were collected in the operating room and processed immediately within 15 min after resection. Snap-frozen tissues were stored at −80°C for subsequent analyses.

#### RT-PCR and sanger sequencing

Total RNA was isolated using TRIzol reagent (Life Technologies, California, USA). First-strand cDNA was synthesized using a High Capacity cDNA Reverse Transcription kit (Life Technologies) according to the manufacturer’s instructions. The amplified bands were gel-purified and later subjected to Sanger sequencing.

#### Quantitative real-time PCR (Qrt-PCR)

cDNA was synthesized from 1.5μg of total RNA using the High Capacity cDNA Reverse Transcription Kit (Life Technologies). Real-time PCR was performed using Power SYBR^®^ Green Master mix (Applied Biosystems, Foster City, USA) and a 7500Fast^™^ Real-Time PCR System (Applied Biosystems). GAPDH gene expression was included as an internal control. The relative expression levels of the fusion genes were calculated using 2^−ΔΔCT^ values. The fusion gene-specific primers and the primers for GAPDH are listed in Supplementary Table 8.

## Acknowledgements

This work was supported by International Science and Technology Cooperation Projects (2015DFA30650 and 2010DFB33720), Capital Special Research Project for Health Development (2014-2-4012), Capital Research Project for the Characteristics Clinical application (Z151100004015170) and Program for New Program for New Century Excellent Talents in University (NCET- 11–0288).

